# MST1R/RON and EGFR in a complex with syndecans sustain carcinoma S-phase progression by preventing p38MAPK activation

**DOI:** 10.1101/252742

**Authors:** DeannaLee M. Beauvais, Kristin Short, Noah Stueven, Scott E. Nelson, Denis Lee, Oisun Jung, Richard A. Anderson, Paul F. Lambert, Alan C. Rapraeger

**Author notes:** Corresponding author: Alan C. Rapraeger, Ph.D. 3053 Wisconsin Institutes for Medical Research University of Wisconsin-Madison 1111 Highland Avenue, Madison, WI 53705.

## Abstract

Syndecan-4 (Sdc4) organizes a complex of receptors consisting of its homologue, Sdc2, the receptor tyrosine kinases EGFR and MST1R/RON, and the laminin-binding α3β1 and α6β4 integrins that depends on a docking site within its extracellular domain. A peptide mimetic of the extracellular docking site, synstatin-EGFR (SSTN_EGFR_), disrupts the receptor complex and prevents the invasion of non-transformed or carcinoma cells that relies on active EGFR. However, the peptide also prevents DNA replication that relies on active MST1R/RON and c-Abl kinase within the complex, resulting in rapid S-phase arrest of head & neck (HN) and breast carcinoma cells. SSTN_EGFR_ does not affect DNA replication in non-transformed oral or breast epithelial cells, but it does block their EGF-dependent invasion. Although EGFR is required as a component of the complex, its kinase activity is not required to sustain S-phase progression in the carcinoma cells, perhaps explaining why many HN and breast carcinomas that overexpress EGFR are nonetheless refractory to EGFR inhibitors. The syndecan-organized receptor complex (Sdc:RTK:ITG complex) appears to suppress stress signals that would otherwise disrupt the replisome engaged in DNA synthesis. SSTN_EGFR_-treatment of carcinoma cells, or normal oral epithelial cells expressing stress-inducing HPV oncogenes, causes rapid activation of the p38 stress MAPK leading to loss of PCNA from the chromatin and cessation of DNA synthesis. This arrest is independent of the common DNA damage response (DDR) known to activate an S-phase checkpoint, revealing a novel arrest mechanism and a novel receptor complex that is activated on tumor cells to suppress stress-induced proliferation arrest.

## INTRODUCTION

The epidermal growth factor receptor (EGFR, HER1) has major roles in human cancer, including squamous cell carcinoma of the head & neck (HNC) and carcinoma of the breast, among others^53, 54, 95^. Traditional causes of HNC are smoking and alcohol consumption, but oropharyngeal tumors caused by human papilloma virus (HPV) infection are increasing dramatically in frequency^101^ and are expected to become the predominant type of cancer caused by HPVs by the end of this decade^22^. EGFR, and its ligands (e.g., EGF, TGFα), are overexpressed in up to 90% of HNC patients, regardless of cause^20, 25, 93^, are induced further by standard treatment regimens involving external beam radiation and DNA damaging agents ^30, 96, 126^, and are strongly linked to tumor progression^21, 87^. EGFR is also a causal agent in triple negative breast cancer, a tamoxifen-resistant, highly malignant form that comprises 15-25% of breast cancers, often striking younger women ^19, 32, 38, 49, 51, 66, 68, 88^. Nonetheless, EGFR inhibitors, including the EGFR-blocking antibody cetuximab or EGFR kinase inhibitors, have had disappointing outcomes in the clinic^20, 48^, suggesting alternative mechanisms through which EGFR promotes or sustains the progression of these cancers.

Work from a variety of laboratories has shown a link between the α6β4 integrin and EGFR in breast and other cancers ^37, 68^. The α6β4 integrin was first identified as a tumor marker for both HNC and breast cancer and has more recently been linked to metastasis and recurrent disease^18, 23, 34, 41, 61, 84, 108, 115, 117^. This integrin in normal cells assembles with laminin in the basement membrane underlying basal epithelial cells as well as endothelial cells lining blood vessels, forming stable hemidesmosomes in which the long (ca. 1000 amino acid) cytoplasmic domain of the β4 subunit anchors to the keratin filament network in the cytoplasm of the cell ^52, 79, 115^. In contrast to this “stabilizing” role, however, the integrin takes part in the invasion, proliferation and survival of tumors that overexpress the receptor tyrosine kinases HER2, EGFR, c-Met and MST1R/RON, including HNC and breast cancer that overexpress EGFR. Signaling from these kinases during wound healing or tumorigenesis causes the breakdown of hemidesmosomes, freeing the integrin to assemble with these kinases ^1, 10, 11, 13, 35, 39, 40, 70, 73, 94, 106, 107, 115^. This leads to its tyrosine phosphorylation that converts it to a signaling scaffold^65, 72, 116^ that binds Shc and IRS-1/2, and activates PI3K, Akt, Erk, Src family members, Rho GTPases and other signaling effectors necessary for proliferation, migration and survival^29, 33, 72, 73, 98, 99, 115, 120,49,50^. Complementing its expression in tumors, EGFR and α6β4 integrin are also expressed in endothelial cells, especially those induced by tumors ^3, 16, 59^. Thus, silencing expression of the β4 subunit or expressing a signaling defective β4 mutant in animal models of breast, prostate or skin cancer dramatically reduces tumorigenesis and/or tumor angiogenesis^27, 44, 105, 122 80^.

Recent work has shown that syndecans act as organizers of receptor complexes critical for cell invasion and survival, particularly in tumor cells ^6–8, 56, 89, 111, 112^. These receptor complexes typically consist of a receptor tyrosine kinase (e.g., IGF-1R, EGFR, HER2, VEGFR2) and an integrin, captured via binding motifs in the syndecan extracellular domains^6–8, 56, 89, 90^. Because these docking sites are extracellular, peptides that mimic these binding motifs (called “synstatins” or “SSTNs”) effectively compete for assembly of the receptor complex and disrupt receptor signaling that is critical for tumor growth and tumor angiogenesis ^5–7, 17, 56, 74–76, 89, 90, 110–114^. This body of work has revealed that the assembly of EGFR with the α6β4 integrin, and with the laminin-binding α3β1 integrin, is mediated by syndecan-4 (Sdc4)^111, 112^. A juxtamembrane site in the extracellular domain of Sdc4 (amino acids 87-131) couples the syndecan to EGFR and the α3β1 integrin^112^, coupling that is competitively blocked by a peptide mimetic called SSTN_EGFR_ ^112^. This peptide blocks the EGF-stimulated migration of keratinocytes and mammary epithelial cells on laminin-332 (LN332, also called LN5) that the cells deposit as they migrate^111, 112^. The C-terminus of the Sdc4 cytoplasmic domain simultaneously captures the α6β4 integrin, an interaction that relies on a glutamate residue (E1729) near the tip of the β4 cytoplasmic domain^111^. Expression of a β4^E1729A^ mutant in keratinocytes and mammary epithelial cells prevents the Sdc4 interaction and also blocks EGF-induced cell migration^111^.

In this study, we have questioned whether Sdc4, EGFR, and the α3β1/α6β4 integrin complex has roles in HNC and breast carcinoma cells other than promoting their invasion. We have used SSTN_EGFR_ to treat carcinoma cells *in vitro* and *in vivo*, and discover that the peptide rapidly induces a novel mechanism of S-phase cell cycle arrest in HNC and breast carcinoma cells. Remarkably, although the arrest mechanism relies on EGFR, it is independent of the EGFR kinase activity and is therefore unaffected by EGFR inhibitors. Instead, sustained progression through S-phase by the tumor cells depends on active macrophage stimulating-1 receptor (MST1R)/Recepteur d’Origine Nantais (RON) kinase ^92, 121^, the Sdc4 homologue Sdc2 ^81^, and the cytoplasmic/nuclear kinase c-Abl ^42^, which are incorporated into the Sdc4-organized receptor complex and are also displaced by SSTN_EGFR_. However, the S-phase arrest is observed only in tumor cells, as the peptide has no effect on the proliferation of non-transformed oral or mammary epithelial cells. However, HPV16 E6 and/or E7 oncogenes, which can immortalize cells but are not sufficient to cause cells to be tumorigenic^50, 58, 118^ convert oral epithelial cells to a SSTN-susceptible phenotype, indicating that preneoplastic and neoplastic carcinoma cells, but not normal epithelial cells, rely on the Sdc4-organized receptor complex to sustain their cell cycle progression in an EGFR kinase-independent manner.

## RESULTS

Our prior work has shown that the survival of SKBr3 breast carcinoma cells depends on a receptor complex in which the EGFR-family kinase HER2 and the laminin-binding α6β4 and α3β1 integrins are coupled by cytoplasmic and extracellular interactions to the matrix receptor Sdc1, contrasting with non-tumorigenic epithelial cells that rely on the receptor complex for migration during wound healing but not for survival^111^. We now question whether a similar tumor-specific mechanism is exhibited by a homologous receptor complex in which HER2 is replaced by the EGFR receptor tyrosine kinase and Sdc1 is replaced by Sdc4 ^111, 112^. The growth of non-tumorigenic oral, skin or breast epithelial cells was compared to a panel of carcinoma cells lines during a four-day treatment with SSTN_EGFR_, a peptide mimetic of the EGFR and α3β1 integrin docking site in the extracellular domain of Sdc4 that competitively displaces these receptors from the syndecan^112^. Whereas the growth of non-tumorigenic normal oral keratinocytes (NOKs), human tonsillar epithelial cells (HTEs), HaCaT keratinocytes and MCF10A cells is not affected (Fig. 1A), HPV^-^ or HPV^+^ HNC cells (Fig. 1B) and triple-negative or HER2+ breast carcinoma cells (Fig. 1C) displayed significantly reduced growth in the presence of the peptide, which exhibited an IC_50_ of approximately 3-10 μM.

**Figure 1.**
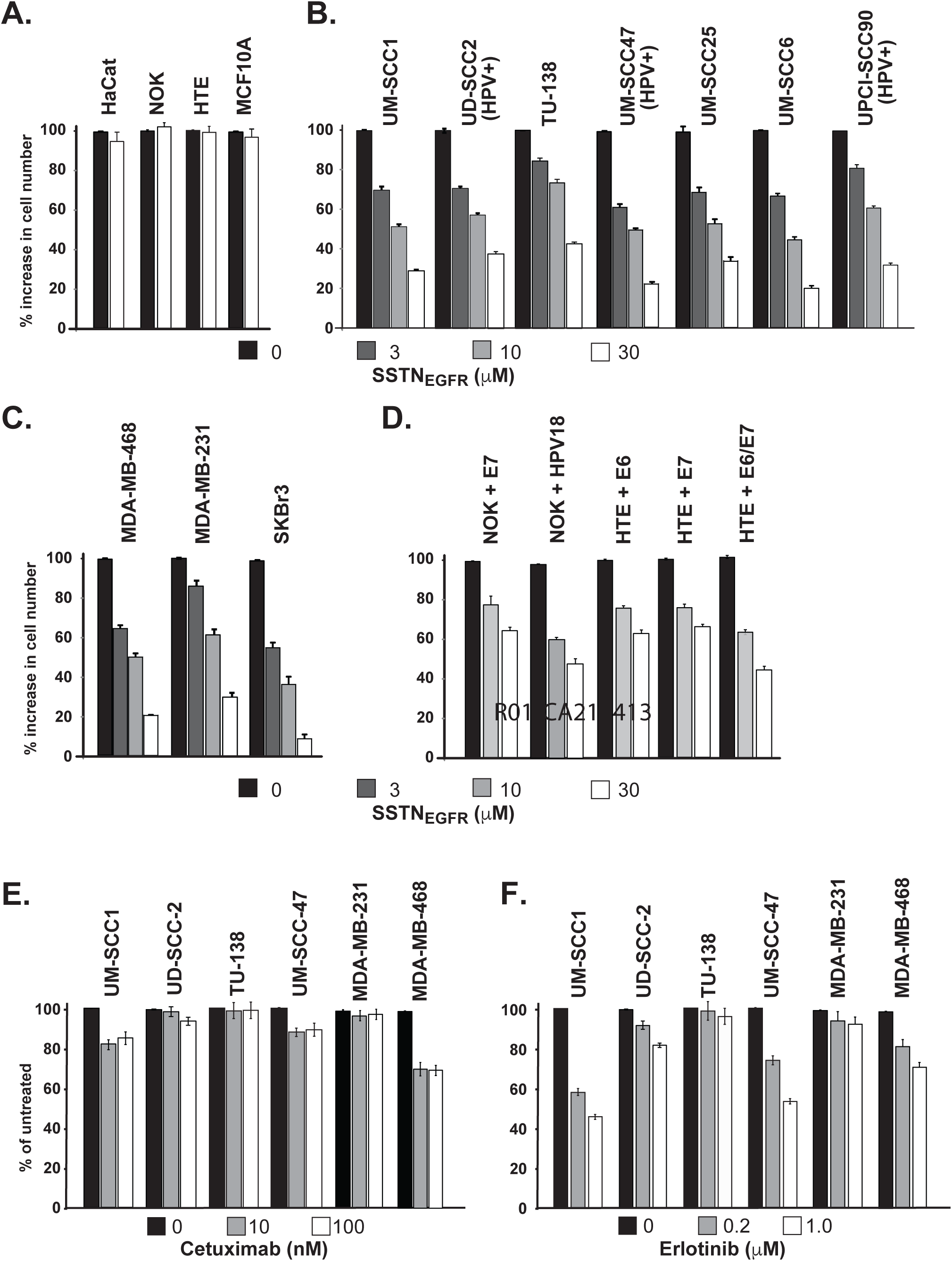
Quantification of impaired growth of SSTN_EGFR_-treated neoplastic and preneoplastic epithelial cells. **A-D**, Cells undergoing logarithmic growth are treated with 0, 3, 10 or 30μ MSSTN_EGFR_ for 4 days, followed by quantification of cell number, expressed as a percentage of untreated (control) cells: **A.** Non-tumorigenic epidermal (HaCaT), oral (NOK, HTE) and breast (MCF10A) epithelial cells; **B.** HPV-negative (UM-SCC1, TU-138, SCC25, UM-SCC6,) and HPV-positive (UD-SCC2, UM-SCC47, UPCI-SCC90) HNC cell lines; **C.** triple-negative (MDA-MB-468, MDA-MB-231) and HER2+ (SKBr3) breast carcinoma; **D.** NOK or HTE cells infected with HPV18 or transfected with HPV16 E6, E7 or E6 and E7 oncogenes (preneoplastic cells); **E, F**. Select HNC and breast carcinoma cell lines are treated with either EGFR-specific inhibitory antibody cetuximab (0, 10 or 100 nM) or kinase inhibitor erlotinib (0, 0.2, or 1.0 μM).

HPV is an increasingly abundant etiological factor in HNC due in large part to the activities of two virally encoded oncogenes E6 and E7, which have been shown to drive head and neck carcinogenesis *in vivo* when mice transgenic for these viral oncogenes are treated with an oral carcinogen^55, 102, 103^. Consistent with these viral oncogenes not being sufficient to cause cancer, human epithelial cells harboring HPV genomes or simply expressing these two oncogenes do not form tumors *in vivo* ^50, 58, 118^. Thus, HPV infection is thought to induce a preneoplastic state, with cancers arising only after the accumulation of additional oncogenic insults. To test whether SSTN targets preneoplastic as well as neoplastic cells, we analyzed the effect of SSTN_EGFR_ on the growth of NOKs or HTEs that were transfected with the HPV16 E6 and/or E7 oncogenes, or infected with HPV18, finding that the introduction of oncogenes into these cells converts them to the SSTN-susceptible phenotype (Fig. 1D).

Overexpression of EGFR and/or its ligands is linked to poor prognosis in HN, lung, triple-negative breast and other carcinomas^20, 48, 51, 66, 88^. Despite this, EGFR inhibitors, including the EGFR-blocking antibody cetuximab, and EGFR kinase inhibitors erlotinib and gefitinib, among others, have had limited success in the clinic ^4, 20, 28, 48^. Similarly, we find that our panel of HNC and triple-negative breast carcinoma cells, despite being highly susceptible to SSTN, are largely resistant to high concentrations of cetuximab (Fig. 1E), and are variably resistant to erlotinib (Fig. 1F). This contrasts with the SSTN_EGFR_ peptide (Fig. 1B, 1C), potentially suggesting a distinct mode of action that depends on EGFR but not EGF ligand or EGFR kinase activity.

Analysis of cell surface receptor expression by flow cytometry demonstrates that all of the cell lines express the component receptors of the Sdc4-organized complex, although the HNC and breast carcinoma cells express 10-20-fold higher levels of EGFR and α6β4 integrin than the non-transformed NOKs or HTEs (Fig. 2A-C). Interestingly, Sdc4 that is responsible for organizing this receptor complex is expressed at nearly undetectable levels in the NOKs and HTEs (Fig. 2B, 2C), cells that appear insensitive to the growth-inhibitory properties of SSTN_EGFR_. However, its levels increase when the NOK or HTE cells express the E6 and/or E7 oncogenes, along with increased expression of EGFR and the α3β1 and α6β4 integrins (Fig. 2B, 2C). This augmented expression of Sdc4 and its presentation at the cell surface is confirmed by western blot analysis of NOK or HTE cells expressing the E6/E7 oncogenes treated with trypsin to remove cell surface Sdc4 prior to cell lysis (Fig. 2D).

**Figure 2.**
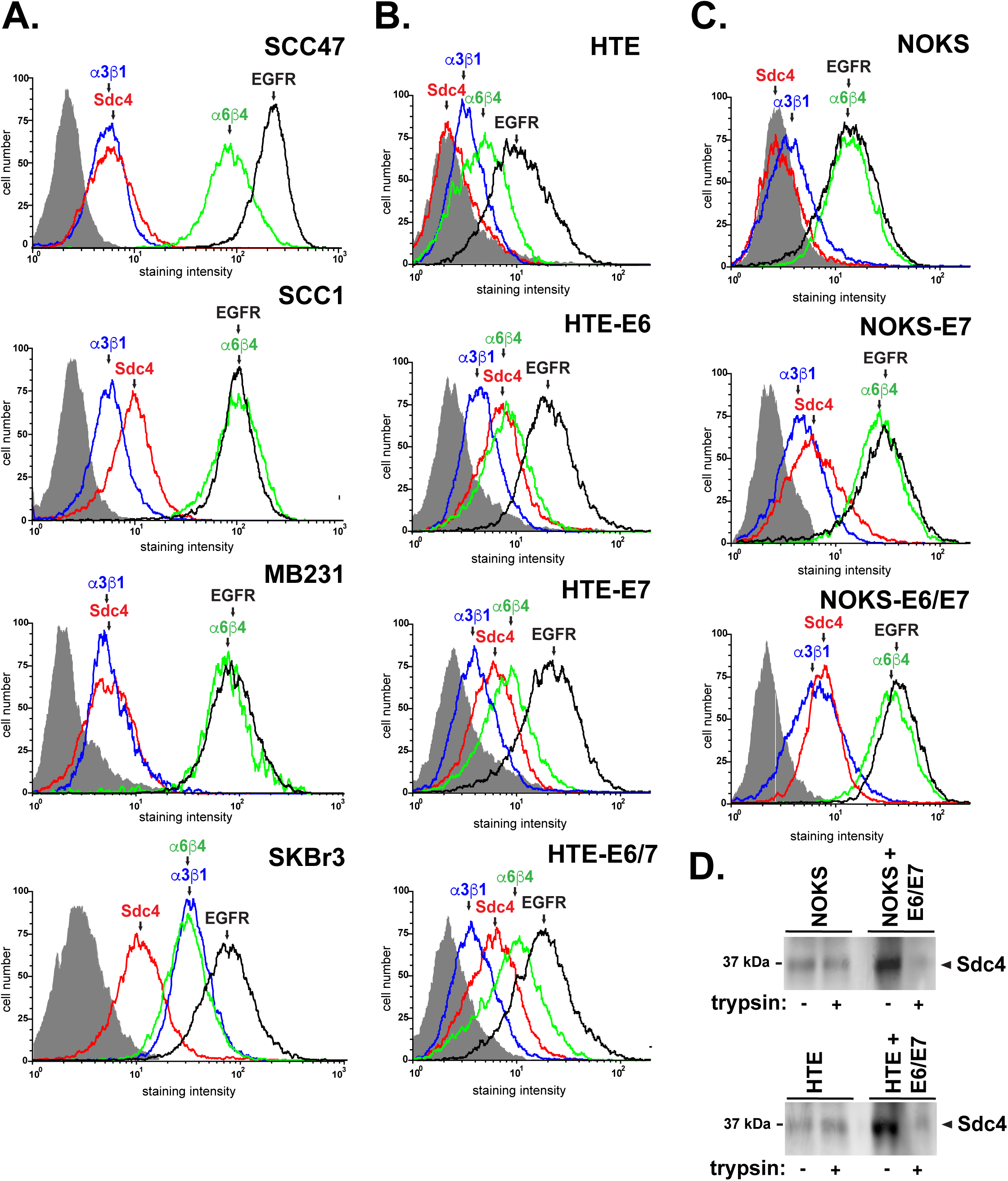
Relative expression of members of the Sdc4:EGFR:ITG complex on non-tumorigenic, preneoplastic and neoplastic epithelial cells. **A-C**, Cell surface of expression of Sdc4 (mAb 8G3, red), α3β1 (mAb P1B5, blue) and α6β4 (mAb3E1, green) integrins, and EGFR (mAb EGFR.1, black) are analyzed by flow cytometry compared to nonspecific IgG (gray profile): **A.** Select HNC (UM-SCC47, UM-SCC1) and breast carcinoma (MDA-MB-231, SKBr3) cell lines; **B.** Non-transformed human tonsillar epithelial cells (HTE) compared to preneoplastic HTE-E6, HTE-E7 cells and HTE-E6/E7 cells; **C.** Non-transformed oral keratinoctyes (NOK) compared to preneoplastic NOK-E7 and NOK-E6/E7 cells. **D.** Western blot analysis of Sdc4 expression in NOKs, NOKs-E6/E7, HTEs or HTEs-E6/E7 whole cell lysates (50 μg total protein/lane) following treatment of the cells with or without trypsin to remove cell surface Sdc4 prior to cell lysis.

The therapeutic potential of peptides is often limited by their rapid destruction by proteases, especially exopeptidases, in the blood plasma and/or their rapid clearance and excretion^109^. However, SSTN_EGFR_ is remarkably stable when incubated at 37°C for prolonged periods in human plasma and displays a half-life of ~42 hr in the plasma of mice following intravenous injection (Fig. 3A, B). The peptide reaches a steady state minimum of 3.2 µM in the plasma with daily injections for 4 days, consistent with this half-life measurement (Fig. 3C). Furthermore, UM-SCC47 xenografts allowed to establish for 1 week in nude mice display ~ 70% reduced growth over the subsequent 4 weeks in the presence of systemic SSTN peptide (Fig. 3D).

**Figure 3.**
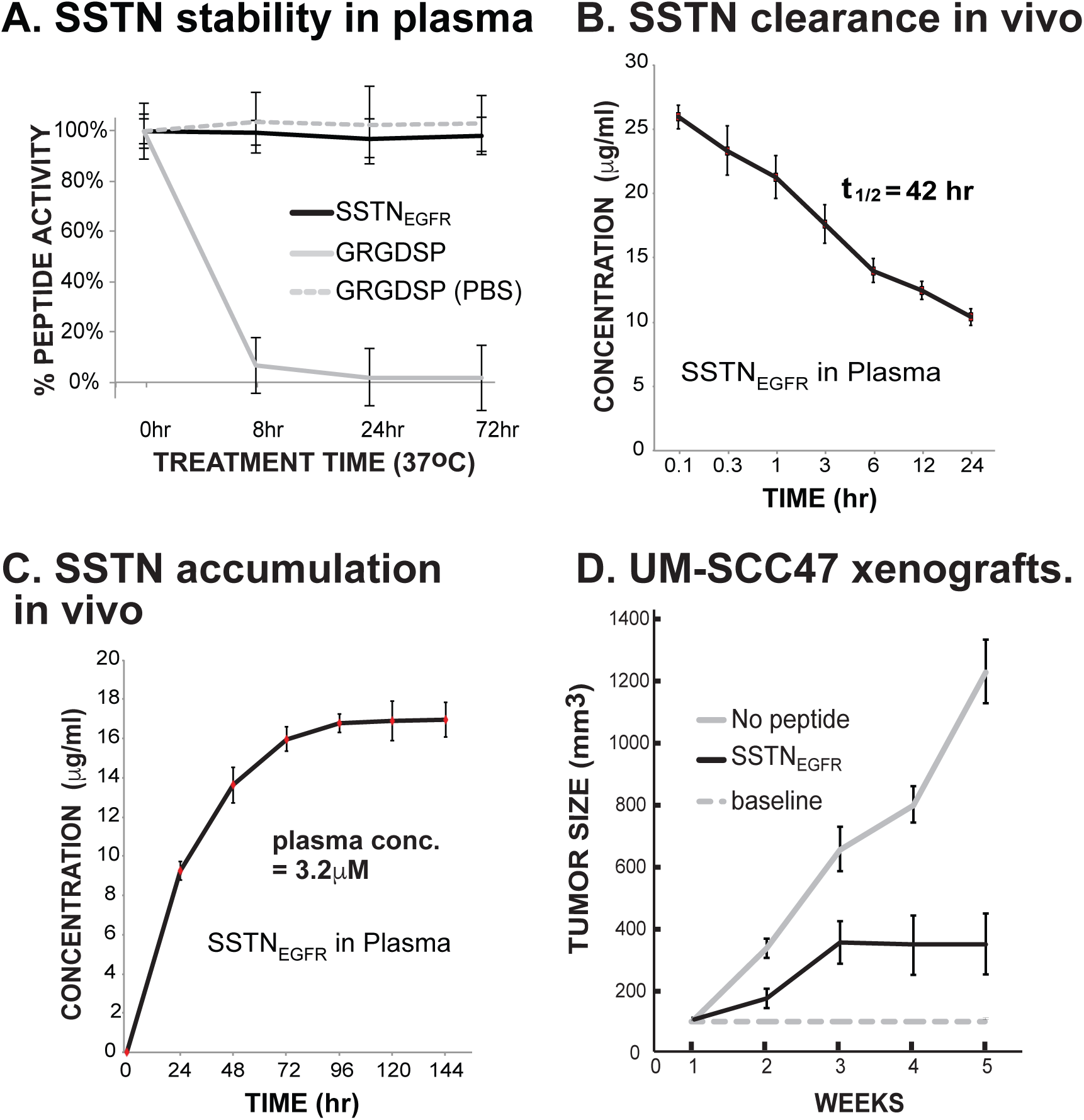
SSTN treatment of UM-SCC47 HNC tumor xenografts. **A.** SSTN_EGFR_, or the GRDGSP integrin inhibitory peptide used as a control, were incubated in either PBS or human plasma for up to 72h at 37°C, then tested for activity. **B.** SSTN half-life following single intravenous (i.v.) injection of 0.597 mg/kg SSTN_EGFR_. **C.** Steady-state SSTN in mouse plasma following daily subcutaneous injection of 0.597 mg/kg peptide. Each data point is the mean +/− SEM of three mice, collected 24 hr after injection. **D.** Subcutaneous UM-SCC47 xenografts in nude mice were treated using Alzet osmotic pump delivery of 1.8 mg/kg/day SSTN_EGFR_ (~10 µM plasma level). Each data point is the mean of six tumors.

While analyzing the response of UM-SCC47 cells to SSTN_EGFR_ *in vitro* by time-lapse microscopy, we observed that none of the treated cells undergo chromosome condensation, form metaphase plates or undergo cytokinesis in the presence of the peptide compared to untreated cells that actively divide to form new daughter cells (data not shown), suggesting that they undergo a cell cycle block prior to reaching M-phase. To examine this possibility, either non-transformed HaCaT keratinocytes or the UM-SCC47 HNC cells line were treated with SSTN_EGFR_ for 3 or 16 hr, accompanied by incorporation of the deoxynucleotide analog 5-ethynyl-2′-deoxyuridine (EdU) into newly synthesized DNA for the last 45 min, and then analyzed by flow cytometry. HaCaT cells treated with the peptide show no significant change in the level of EdU incorporation into cells nor a change in the distribution of cells in G1, S or G2/M phases of the cell cycle (Fig. 4A). In contrast, the UM-SCC47 cells show dramatically reduced EdU incorporation already by 3 hr of treatment (Fig. 4A). Moreover, the percentage of cells in G2/M is dramatically reduced in agreement with our time lapse microscopy observation, whereas the percentage of cells in S-phase or G1 is relatively unchanged (Fig. 4A). However, it is possible that many of the cells with 2N content that are assigned to G1 are actually arrested in very early S-phase, especially after 16 hr of treatment, suggesting that carcinoma cells treated with the peptide when in early-, mid- or late-S-phase undergo a rapid arrest of DNA synthesis. To test this hypothesis, UM-SCC47 cells were subjected to a double-thymidine block to synchronize them at the G1/S interface, then released for 3 or 8 hr to proceed either into early or late S-phase prior to being treated with SSTN (Fig. 4B). Cells released for 3 hr, then chased for an additional 8 hr with vehicle alone appear to progress to late S-phase/G2/M, or pass through G2/M and enter G1, as shown by their 2N DNA content (Fig. 4B). However, cells released for 3 hr then chased for 8 hr with SSTN_EGFR_ remain almost entirely in S-phase and show little or no EdU incorporation above background. Similarly, after an 8 hr release, the cells appear to be in late S-phase, and proceed almost totally to G1 when treated with vehicle alone for an additional 8 hr, whereas over half of the cells remain arrested in late S-phase if chased in SSTN_EGFR_ (Fig. 4B). The remainder enter G1, suggesting that peptide-mediated arrest is not immediate and those cells nearing the end of S-phase at the time of peptide addition continue into G2/M and are no longer subject to arrest by the peptide.

**Figure 4.**
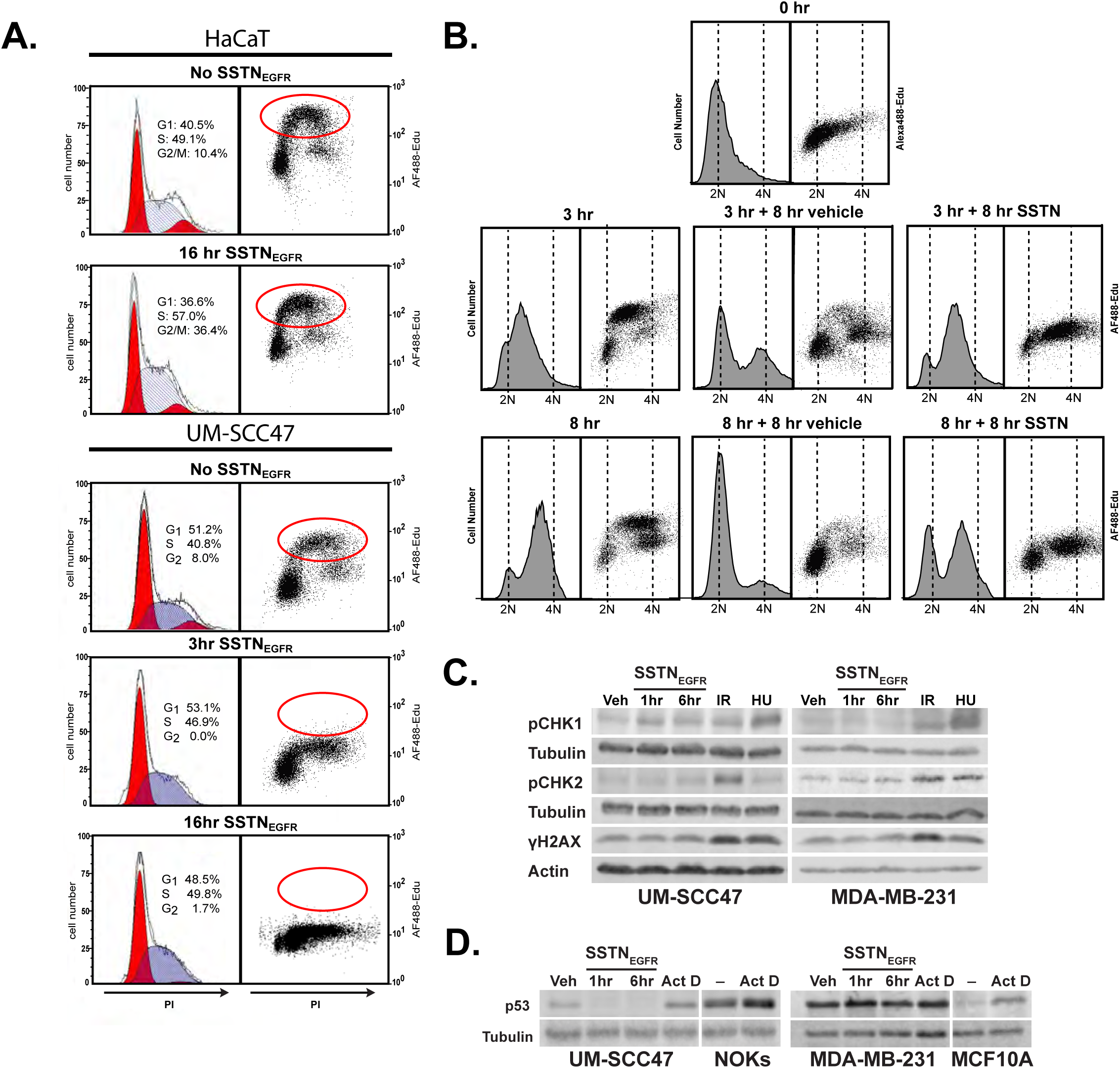
SSTN_EGFR_ causes S-phase arrest of neoplastic and preneoplastic epithelial cells. **A.** HaCaT keratinocytes or UM-SCC47 cells were treated with or without 30 uM SSTN_EGFR_ for up to 16 hr. EdU was added for the last 45 min, followed by PFA fixation, saponin permeabilization, staining with Alexa488 azide (Click-IT labelling reaction) and propidium iodide (PI), and flow cytometry analysis to monitor EdU incorporation and DNA content, respectively; **B.** UM-SCC47 cells are arrest in early S-phase using a double thymidine block, released for either 3 or 8 hr, then subjection to 8 hr treatment with either 30 µM SSTN_EGFR_ or vehicle alone. EdU was added for the last 45 min and the cells were analyzed by flow for DNA content and EdU incorporation. **C.** UM-SCC47 or MDA-MB-231 cells were treated with either vehicle alone for 6 hr, 30 μM SSTN_EGFR_ for either 1 or 6 hr, 8 Gy ionizing radiation (IR), or 5 mM hydroxyurea (HU) for 1 hr prior to lysing and analysis for activation of DNA damage response effectors using mAbs to pS345-Chk1, pT68-Chk1 or pS139-H2AX. **D.** UM-SCC47 and MDA-MB-231 cells were treated with vehicle alone for 6hr, 30 μM SSTN_EGFR_ for 1 or 6 hr, or 5 nM Actinomycin D (Act D) for 24 hr to induce stress, then lysed and analyzed for total p53 protein. Non-transformed NOK or MCF10A cells were treated with or without Act D as positive controls for the stress response. Tubulin (**C and D**) and actin (**C**) serve as loading controls.

Growth factor receptor signaling is typically required to bypass the G1/S “start” point of the cell cycle, but continued growth signaling is not required to sustain cell cycle progression once DNA synthesis has begun^83^. Mechanisms that do cause arrest at the S-phase checkpoint arise from activation of the DNA damage response (DDR), due to actual DNA damage or replicative stress^36, 57^. This activates p53 and/or checkpoint kinases (e.g. Chk1 or Chk2) downstream of the DNA damage sensors ATM, ATR and DNA-PK leading to phosphorylation of regulatory factors, including the histone H2AX^14, 36, 57^. However, treatment of UM-SCC47 HNC or MDA-MB-231 breast carcinoma cells with SSTN even for 6 hr fails to activate Chk1 or Chk2, or cause phosphorylation of H2AX (γH2AX), in contrast to the phosphorylation observed then the cells are treated with external beam radiation to induce DNA breaks or hydroxyurea to induce replicative stress that are known to activate the DDR (Fig. 4C)^14^. Similarly, the UM-SCC47 cells, which express wild-type p53^15^, appear to lose rather than stabilize p53 when treated with SSTN, in contrast with the induction of p53 by Actinomycin D-induced stress (Fig. 4D). MDA-MB-231 cells, which express abundant levels of mutant p53 that escapes MDM2-mediated degradation^82^, fail to alter p53 levels in response to either SSTN or Actinomycin D (Fig. 4D).

These findings suggest that the Sdc4:ITG:EGFR complex has a prominent role in the progression of preneoplastic and neoplastic epithelial cells through S-phase. To confirm this, we compared EdU incorporation in non-tumorigenic cells (NOKs and MCF10A), preneoplastic cells (NOKs expressing E6/E7 oncogenes) and either HN or breast carcinoma cells treated with SSTN_EGFR_ for 1, 3, or 6 hr. As expected (cf. Fig. 1) the NOKs and MCF10A cells show no inhibition of DNA synthesis at any treatment time, including as long as 16 hr (data not shown), whereas the preneoplastic and neoplastic cells show a dramatic reduction in EdU incorporation even when pretreated with SSTN_EGFR_ for as little as 1 hr. Note that EdU is added together with continued SSTN for 45 min following the initial 1, 3 or 6 hr pretreatment with SSTN (Fig. 5A).

**Figure 5.**
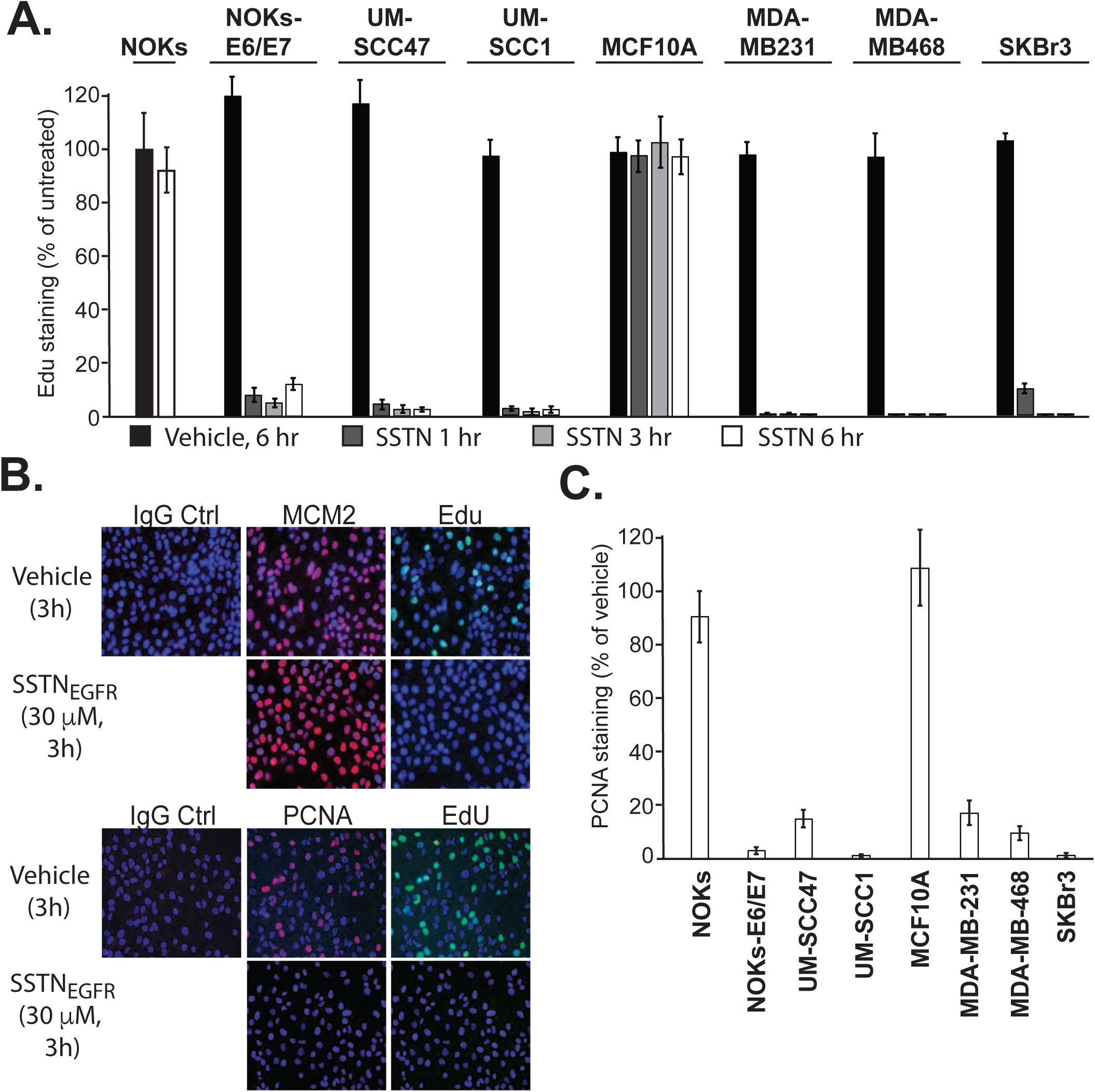
Diminished DNA synthesis is mirrored by loss of PCNA. **A.** Select neoplastic, preneoplastic and non-transformed epithelial cells were cultured with or without 30 μM SSTN_EGFR_ for up to 6 hr, followed by EdU labeling for 45 min in the continued presence of the peptide. EdU incorporation is expressed as a percentage of that observed in cells grown in the presence of vehicle alone; **B.** UM-SCC47 cells were treated with 30 μM SSTN_EGFR_ or vehicle for 3 hr, with EdU added for the last 45 min, then extracted and stained to detect EdU incorporation and together with either DNA-bound PCNA or the origin licensing factor MCM2; **C.** Quantification of PCNA staining in HN or mammary cells treated either with vehicle or 30 μM SSTN_EGFR_ for 3 hr.

A key feature of DNA synthesis is assembly of the replisome, organized by the trimeric PCNA sliding clamp that encircles the DNA strand and acts as a scaffold to assemble DNA polymerase and its regulatory proteins on the DNA^69, 78^. PCNA loading occurs on thousands of sites that have been licensed throughout the genome during G1, including the licensing factors ORC, Cdc6, Cdt1, and the MCM2-7 complex^12^. To test whether the loading of PCNA or the assembly of the licensing complex depends on a signal from the Sdc4-organized receptor complex, UM-SCC47 cells were detergent-extracted following 3 hr treatment with SSTN_EGFR_ and stained for the retention of MCM2 and PCNA on the insoluble chromatin fraction (Fig. 5B). As expected, SSTN causes the near complete loss of EdU incorporation (Fig. 5B). This is mirrored by loss of PCNA as well (Fig. 5B), whereas MCM2 staining appears unaffected. Similar staining is observed for ORC (data not shown), suggesting that PCNA loading/unloading is the target of the inhibitory signal rather than the licensing complex. PCNA loss in the presence of SSTN extends to triple negative and HER2+ breast carcinoma cells, select HPV- (UM-SCC1) and HPV+ (UM-SCC47) HNC cells and preneoplastic NOKs expressing E6/E7 oncogenes (Fig. 5C), but not to non-transformed NOKs or MCF10A mammary epithelial cells (Fig. 5C), mirroring the block to EdU incorporation (Fig. 5A).

To examine which of the receptors in the Sdc4:ITG:EGFR complex is responsible for signaling that sustains the carcinoma S-phase progression, each of the component receptors was silencing in UM-SCC47 cells using transient (72 hr) siRNA treatment, resulting in >95% reduction in receptor expression (Fig. 6A). Regardless of which receptor was silenced (.g., Sdc4, integrin α3β1, integrin α6β4 or EGFR), DNA synthesis was blocked by well over 90% (Fig. 6A).

**Figure 6.**
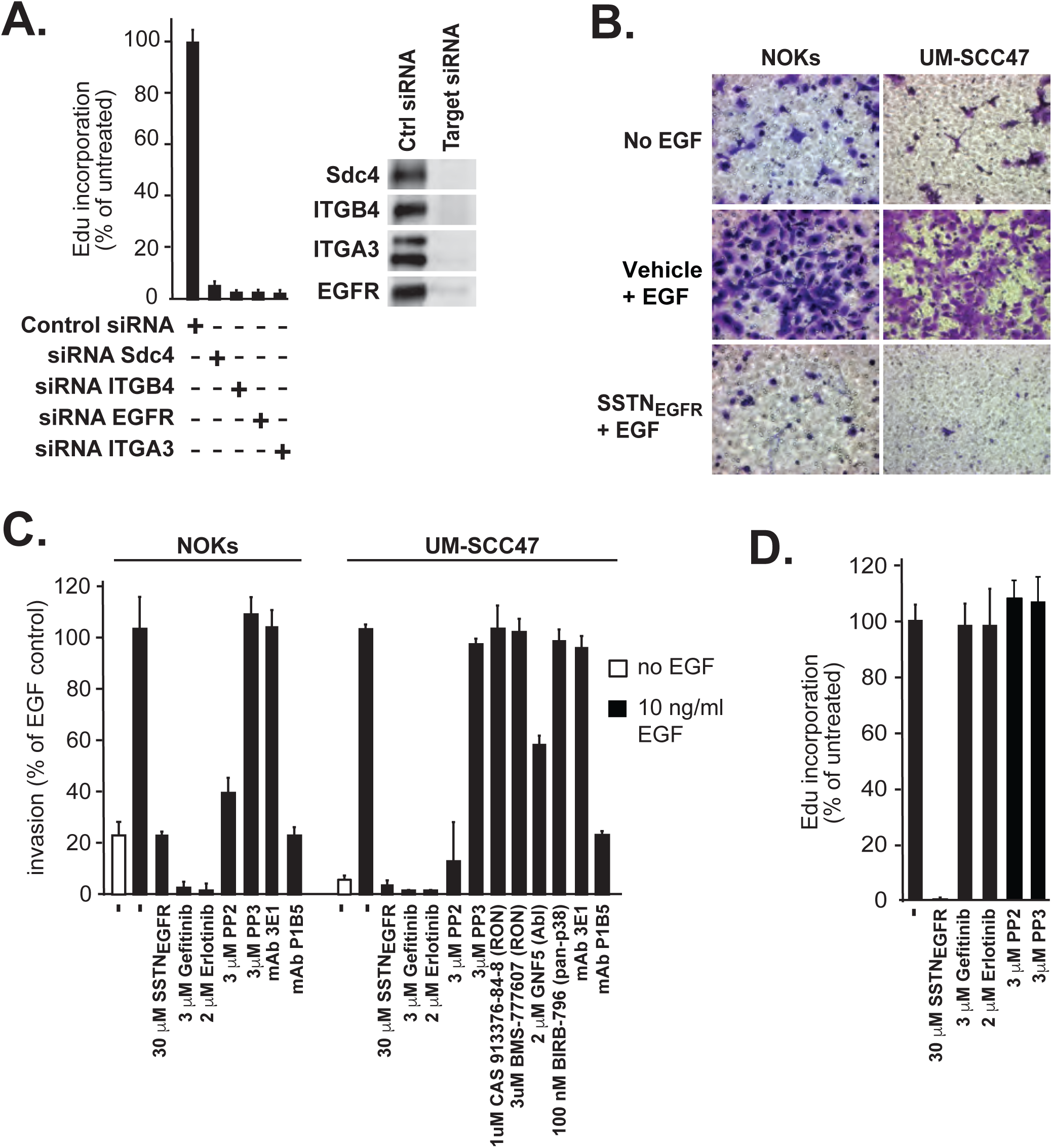
Role of the individual receptors in cell invasion or S-phase progression. **A.** UM-SCC47 cells were treated for 72 hr with either control siRNA, or siRNA specific for human Sdc4, β4 integrin (ITGB4), EGFR, or α3 integrin (ITGA3), followed by addition of EdU for 45 min to monitor DNA synthesis prior to fixation and staining; Western blots showing individual receptor expression 72 hr after siRNA transfection are shown (inset); **B**. Either NOKs or UM-SCC47 cells were allowed to invade through LN332-coated filters for 16 hr in serum-free conditions following stimulation with 10 ng/ml EGF +/− 30 μM SSTN_EGFR_ or vehicle; **C.** Quantification of NOK or UM-SCC47 cell invasion as in (**B**) in the presence or absence of the EGFR kinase inhibitors gefitinib (3 μM) or erlotinib (2 μM), SFK inhibitor PP2 or its inactive analogue PP3 (3 μM), MST1R/RON inhibitors CAS 913376-84-8 (1 μM) or BMS-0777607 (3 μM), c-Abl inhibitor GNF5 (2 μM), pan-p38MAPK inhibitor BIRB-796 (100 nM) and α6β4 (3E1) or α3β1 (P1B5) integrin blocking antibody (10 μg/ml); **D.** DNA synthesis is detected by EdU incorporation in UM-SCC47 cells grown for 3 hr in medium containing SSTN_EGFR_, the EGFR kinase inhibitors gefitinib or erlotinib, or SFK inhibitor PP2 versus its inactive analogue PP3.

We have shown previously that EGFR and the α3β1 and α6β4 integrin coupled to Sdc4 play a prominent role in the LN332-dependent migration of epithelial cells ^111, 112^, which depends on stimulation with EGF to activate the EGFR, resulting in activation of the Src-family kinase Fyn and tyrosine phosphorylation of the long cytoplasmic tail of the α6β4 integrin tethered to the cytoplasmic domain of Sdc4 ^29, 73^. Revisiting this role in the context of oral epithelial and carcinoma cells, both NOKs and UM-SCC47 cells are observed to invade through LN332-coated filters in response to EGF, with little to no invasion observed under serum-free conditions if EGF is not added (Fig. 6B, 6C), Furthermore, SSTN_EGFR_, the EGFR kinase inhibitors erlotinib or gefitinib, or the Src-family kinase (SFK) inhibitor PP2 reduce cell invasion either to or below the levels observed when EGF is omitted (Fig. 6C). Invasion is also blocked by the α3β1 integrin blocking antibody P1B5, which prevents the integrin’s interaction with LN332^111^. Although α6β4 integrin is also a LN332-binding integrin, mAb 3E1 that blocks this integrin does not block invasion, confirming our prior findings that although the presence of this integrin is required, its primary role is as a signaling component/scaffold and it’s engagement with the matrix is not necessary^111, 112^. A series of other inhibitors that target MST1R/ RON, a c-Met family member^92, 121^, or p38MAPK, have no effect, whereas inhibition of the cytoplasmic/nuclear kinase c-Abl, known to be activated downstream of integrins^63, 64^, had a modest inhibitory effect (Fig. 6C).

To test whether kinase activation responsible for cell invasion is also required for tumor cell advancement through S-phase, UM-SCC47 cells were labeled with EdU while being treated with erlotinib, gefitinib, or PP2. Surprisingly, despite the loss of cell proliferation when EGFR expression is silenced (Fig. 6A), neither inhibition of EGFR or SFKs prevent EdU incorporation in the UM-SCC47 cells (Fig. 6D). This prompted us to question the potential role of other kinases. MST1R/RON has been reported previously to associate with the α6β4 integrin^94, 123^, and also to activate c-Abl in prostate cancer cells leading to tyrosine phosphorylation of PCNA^124, 125^. Using specific inhibitors to MST1R/RON or Abl kinase, we find that EdU incorporation (Fig. 7A) and PCNA retention on chromatin (Fig. 7B) is dramatically reduced in the preneoplastic NOK-E6/E7 cells and the HN and breast carcinoma cells, but not in the NOKs or MCF10A cells, duplicating what we find with SSTN_EGFR_ treatment. This prompted us to test whether Sdc4 organizes additional receptors into the complex with EGFR. Indeed, probing Sdc4 immunoprecipitates from UM-SCC47 cells reveals that MST1R/RON, c-Abl and the Sdc4 homologue, Sdc2^81^ also precipitate (Fig. 7C). MST1R/RON and c-Abl are active, as evidenced by pY1238/1239 and pY412 in the A-loops of their kinase domains, respectively, and pY245 in the SH2-kinase linker of c-Abl^47, 121^, suggesting that they are activated by their incorporation into the complex. The SSTN_EGFR_ peptide, which mimics the extracellular docking site unique to Sdc4, displaces MST1R/RON, Abl kinase and Sdc2 from the complex, along with EGFR and the α3β1 integrin (Fig. 7C). The α6β4 integrin remains due to its docking to Sdc4 by its cytoplasmic domain^111^. Furthermore, the displacement of MST1R/RON and c-Abl by SSTN_EGFR_ inactivates c-Abl (Fig. 7D), supporting the hypothesis that their incorporation into the complex causes their activation and generates the signaling that sustains S-phase progression.

**Figure 7.**
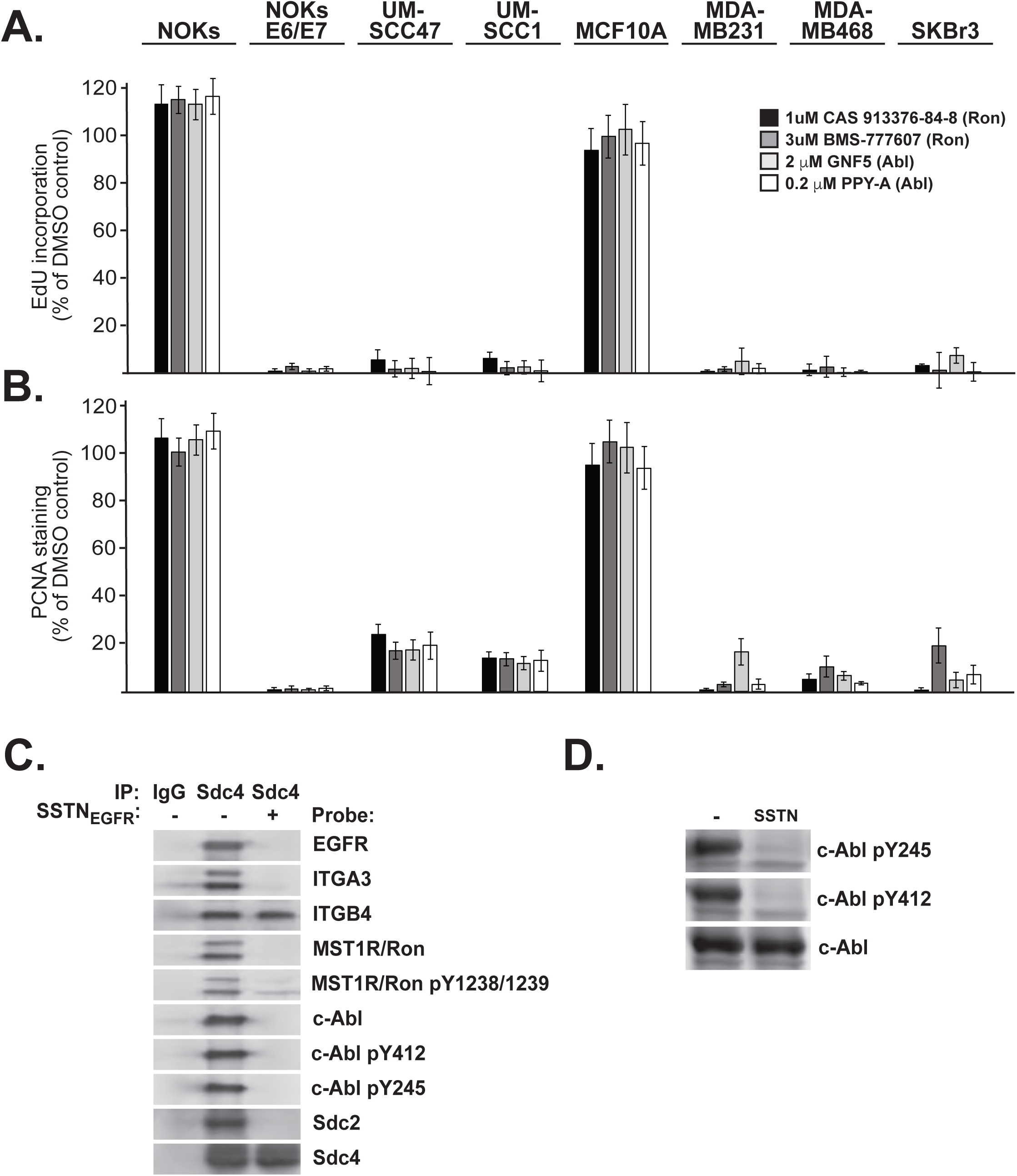
Active MST1R/RON and Abl kinases are required to sustain S-phase progression in carcinoma cells. **A, B.** Select neoplastic, preneoplastic and non-transformed epithelial cells were treated for 3 hr in DMSO (vehicle), the MST1R/RON inhibitors CAS 913376-84-8 (1 μM) or BMS-777607 (3 μM) or the Abl kinase inhibitors GNF5 (2 μM) or PPY-A (0.2 μM). EdU was added for the last 45 min. The cells were fixed and extracted to stain for EdU incorporation (**A**) or DNA-bound PCNA (**B**); **C.** UM-SCC47 cells treated with vehicle alone or 30 μM SSTN_EGFR_ for 3 hr were lysed and subjected to immunoprecipitation with non-specific isotype control IgG or mAb 8G3 to Sdc4. Immunoprecipitates were probed for the presence of EGFR, α3 integrin (ITGA3), β4 integrin (ITGB4), total and active MST1R/RON (pY1238/1239), total and active c-Abl (pY412 and pY245), Sdc2 and Sdc4; **D**. UM-SCC47 cells were treated for 3 hr with vehicle or 30 μM SSTN_EGFR_, then extracted and analyzed on western blots for active pY245 or pY412 phosphorylated c-Abl.

Neoplastic cells are under oncogenic, oxidative, metabolic and genotoxic stress that sets them apart from non-tumorigenic cells^14, 45, 46, 71^. Our prior work has shown that yet another syndecan-organized receptor complex consisting of IGF-1R and the αvβ3 or αvβ5 integrin coupled via extracellular domain interactions to Sdc1 serves in tumor cells to suppress the activation of apoptosis signal-regulating kinase-1 (ASK1), a MAP3K responsible for the activation of Jun N-terminal kinase (JNK) leading to apoptosis^8^. To test the hypothesis that the Sdc4-organized receptor complex serves in a similar manner to suppress stress signals that might otherwise cause S-phase arrest, we probed an antibody array consisting of apoptotic markers with lysates of cells treated for 16 hr with SSTN_EGFR_. Among the changes noted in UM-SCC47 and SCC25 HNC cells, but not non-transformed HaCaT, NOK or MCF10A cells, was an upregulation of active p38 stress MAPK (p38MAPK) (Fig. 8A). Blot analysis of oral epithelia over a series of SSTN treatment time points ranging from 15 min to 16 hr confirm that the NOKs fail to activate p38MAPK in response to SSTN_EGFR_, but NOKs expressing the E6/E7 oncogenes and HNC cells activate p38MAPK within 15 min of peptide addition, and the activation persists throughout the period examined (Fig. 8B, 8C). A similar finding is observed in breast carcinoma cells, in which p38MAPK is activated within 15 min in tumor cells but no activation is witnessed in non-tumorigenic MCF10A cells (Fig. 8D). Furthermore, UM-SCC47 HNC cells treated with SSTN_EGFR_ in the presence of p38MAPK inhibitors, BIRB-796 (Doramapimod) or Losmapimod (GW856553X, SB856553, GSK-AHAB) - at concentrations sufficient to block its activation (Fig. 8F) - fail to arrest their DNA synthesis, as shown by EdU staining (Fig. 8E). This extends as well to each of the preneoplastic or neoplastic oral and breast cells, in which addition of p38MAPK inhibitors fully rescues the inhibitory effect of SSTN_EGFR_ on DNA synthesis based on EdU incorporation (Fig. 8G).

**Figure 8.**
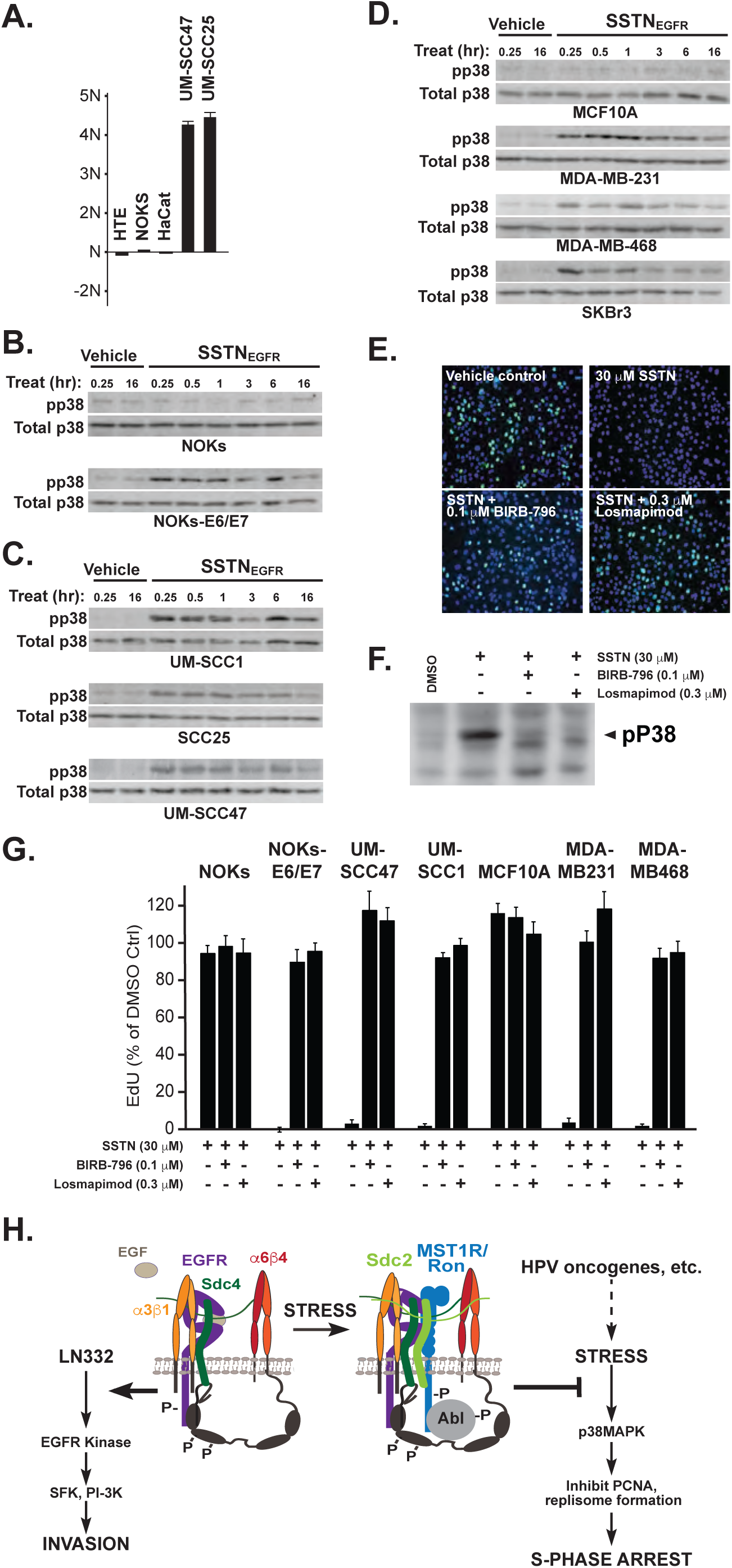
SSTN-induced cell cycle arrest depends on activated p38MAPK. **A.** Quantification of p38MAPK activation by antibody array (pT180/pY182) in non-transformed (HTE, NOK, HaCaT) keratinocytes and HNC cells (UM-SCC-47, UM-SCC25). **B-D**, Detection of pT180/pY182 p38MAPK following treatment with vehicle alone or 30 μM SSTN_EGFR_ in NOKs or NOKs expressing the HPV16 E6/E7 oncogenes (**B**), in HNC cells (**C**), or **in** non-transformed (MCF10A) or transformed (MDA-MB-231, MDA-MB-468, SKBr3) mammary epithelial cells (**D**); **E.** UM-SCC47 cells treated with 30 μM SSTN_EGFR_ in the presence or absence of p38MAPK inhibitors 0.1 μM BIRB-796 or 0.3 μM Losmapimod for 1 h followed by continued treatment for 45 min in the presence of EdU; **F**. Western blot analysis of p38MAPK inhibition by 0.1 μM BIRB-796 or 0.3μM Losmapimod in UM-SCC47 cells (30 μg total protein/lane); **G.** Quantification of DNA-bound PCNA in select cells treated with 30 μM SSTN_EGFR_ with or without p38MAPK inhibitors for 3 hr; **H**. Model showing role of the Sdc:RTK:ITG complex in promoting EGFR kinase-dependent cell invasion and its EGFR kinase-independent suppression of stress-induced p38MAPK activation, thereby allowing normal PCNA loading/unloading in tumor cells.

## DISCUSSION

We have defined a novel signaling mechanism that sustains the proliferation of preneoplastic epithelial cells and HN and breast carcinoma cells. This mechanism builds on a previously described signaling complex organized by Sdc4 at the surface of migrating epithelial cells, comprised of the α3β1 integrin and EGFR docked to a juxtamembrane site in the Sdc4 extracellular domain, and the α6β4 integrin engaged with the Sdc4 cytoplasmic domain following its release from hemidesmosomes during the initiating steps of cell migration ^111, 112^. Migration depends on active EGFR kinase and Fyn-mediated phosphorylation of the β4 integrin cytoplasmic domain, engaging signaling effectors that drive migration^29, 33, 72, 73, 98, 99, 115, 120,49,50^. We now show that additional components, namely, MST1R/RON, c-Abl and Sdc2 are components of the complex as well, although engaged in a different role. Whereas they have little or no role in cell invasion, they are critical for sustaining the proliferation of tumor cells or epithelial cells expressing oncogenes that put the cells under stress. The role of Sdc2 remains unexplored, but it may play in important role by bringing these new receptors to the complex, facilitated by its heterodimerization with Sdc4^24^. MST1R/RON has also been shown to form heterodimers with EGFR^60, 85^, potentially further stabilizing its incorporation into the receptor complex. When incorporated, both MST1R/RON and c-Abl are activated, providing the signaling necessary to suppress stress signaling. By carrying out this suppression, the syndecan-organized receptor complex prevents p38MAPK from phosphorylating unknown targets that cause the collapse of the replisome and cessation of DNA synthesis resulting in a novel S-phase arrest.

Cellular stress can cause arrest in all phases of the cell cycle. DNA damage or stressors that arrest replication (e.g., hydroxyurea) trigger the DDR, which activates ATM and ATR^26, 36^ and their effector kinases Chk1 (ATR) and Chk2 (ATM). These kinases arrest DNA synthesis in S-phase - at least in part - by targeting the cdc25A phosphatase and preventing its activation of cyclin E/Cdk2 required to bypass the G1/S checkpoint, and cyclin A/Cdk2 necessary for DNA synthesis in S-phase^100, 119^. Also, Chk1 phosphorylates p21^WAF1/CIP1^ to disrupt cyclin E/Cdk2, ATR reverses the degradation of p16^INK4A^, inhibiting cyclin D/Cdk4/6 and its phosphorylation of retinoblastoma protein (RB), arresting cells in G_1_ ^2^. Chk2 targets CDC25B phosphatase, preventing its activation of cyclin B/Cdk2 and causing arrest in G_2_/M. These kinases also target p53, affecting both its stability and transcriptional activity^86^, leading to G_1_ and G_2_/M arrest through its control of p21 and cdc25A phosphatase expression. Despite the utilization of this mechanism to activate the S-phase checkpoint by a wide variety of stressed or damaged cells, however, the S-phase arrest induced by disruption of the syndecan:receptor tyrosine kinase:integrin (Sdc:RTK:ITG) signaling complex appears to be distinct from the canonical DDR, as treatment with SSTN_EGFR_ fails to activate Chk1, Chk2, or p53.

Increasing evidence suggests that p38MAPK and JNK stress kinases have significant roles in cell cycle arrest, whether caused by DNA damage or other types of stress (e.g., oncogenic stress, metabolic stress)^43, 67, 97^. Targets of p38MAPK and/or JNK include: (i) the aforementioned stabilization of p53, (ii) MAPK-activated protein kinase 2 (MAPK-APK2 or MK2) that targets and inactivates the cdc25A phosphatase, thus inducing G1 and S-phase arrest^91^, and (iii) phosphorylation of Cdt1, a critical licensing factor that, together with cdc6 and the histone deacetylase HB01, mediates the assembly of the minichromosome maintenance complex (MCM) at origins of replication during G1, thus completing origin licensing necessary for cells to progress into S-phase^9, 43, 77^. In contrast to these outcomes, however, the rapid, p38MAPK-dependent loss of PCNA from sites of replication when tumor cells are treated with SSTN_EGFR_, despite the apparent retention of MCM2 as a marker for of the licensing complex to which it binds, suggests that the p38MAPK target is part of the replisome. PCNA is a homotrimeric sliding clamp that acts as a scaffold for the replication machinery, including DNA polymerase δ and ɛ, cyclin A/CDK2, GADD45, p21^CIP1/WAF1^, and other regulatory proteins^69, 78, 62^. The loading/unloading of PCNA is regulated by the replication factor-C complex (RF-C); the largest of its five subunits, RF-Cp145, is phosphorylated by CDK2 when activated by cyclins, especially cyclin B in G2/M that causes release of PCNA from the DNA at the culmination of S-phase. But, whether it could also be a target of p38MAPK during S-phase is not known. Intriguingly, phosphoproteomic screening indicates that PCNA itself is phosphorylated on S72 (FFRL**S**PKDSE)^31^ in a consensus site potentially recognized by stress MAPKs and an area of future examination.

Normal oral and breast epithelial (e.g., non-transformed) cells are refractory to SSTN-induced S-phase arrest, whereas they become susceptible if expressing HPV oncogenes. They also upregulate expression of the complex components, especially the organizer, Sdc4, suggesting a feedback mechanism in response to the oncogenes. We suggest that this is a response to cellular stress in response to E6 and E7 expression (see model, Fig. 8H). Another response to this stress is activation of p38MAPK, destined to arrest the cells in S-phase by displacing PCNA from the DNA, a response that is prevented by the upregulated Sdc:RTK:ITG complex. The fact that non-transformed epithelial cells fail to arrest is likely due to the fact that they are not under the sustained stress that transformed cells experience.

The signaling mechanism that suppresses the stress signal appears to depend on MST1R/RON and c-Abl. MST1R/RON, and its family member c-Met, have been shown to associate with the α6β4 integrin^94, 123^, although whether the interaction is direct or through other receptors such as the complex described here is not clear. Because inhibition of either MST1R/RON or c-Abl causes the S-phase arrest, it is likely that c-Abl is activated by MST1R/RON, as has been shown in other cancers^125^. However, the target of c-Abl that restricts p38MAPK activation, presumably an upstream enzyme, remains unknown. It is interesting to note that PCNA itself is phosphorylated on Y211 by c-Abl^124^, and reportedly by nuclear EGFR as well, protecting PCNA from proteasome-mediated degradation^56, 78, 79, 88^. Although blockade of such phosphorylation when c-Abl activation is blocked by SSTN_EGFR_ is unlikely to explain p38MAPK-mediated S-phase arrest that we observe, it may well be an additional outcome of c-Abl activation by the Sdc:RTK:ITG complex.

## MATERIALS AND METHODS

### Reagents

SSTN_EGFR_ peptide obtained from LifeTEIN LLC (Hillsborough, NJ) was reconstituted in DMEM (Life Technologies, Grand Island, NY) containing 50 mM HEPES (Millipore-Sigma, Burlington, MA) for *in vitro* studies, or HEPES-buffered 0.9% saline for use *in vivo*.

Monoclonal antibody (mAb) against human Sdc4 (F94-8G3) was graciously provided by Drs. Guido David and Pascale Zimmerman (University of Leuven, Belgium and Centre de Recherche en Cancérologie de Marseille, France), integrin α6β4 (3E1) was obtained from the Memorial Sloan-Kettering Monoclonal Antibody Core Facility (New York, NY) and integrin α3β1 (P1B5) from Millipore-Sigma. Mouse mIgG1 and mIgG2b isotype controls and mAb EGFR.1 were from BD Biosciences (San Jose, CA). Western blotting primary antibodies include: goat polyclonal human Sdc4 (AF2918, 0.5 μg/mL), EGFR (AF231, 1 μg/mL) and total MST1R/RON kinase (AF691, 1 μg/mL); rabbit polyclonal pY1238/1239 MST1R/RON (AF1947, 1 μg/mL) and mouse human ITGB4 mAb 422325 (1 μg/mL) from R&D Systems (Minneapolis, MN); rabbit polyclonal ITGA3 (1:250) from Novus Biologicals (Littleton, CO); rabbit mAbs 73E5 (c-Abl pY245, 1 μg/mL), 247C7 (c-Abl pY412, 1 μg/mL), D13E1 (total p38MAPK, 1:1000), 11H10 (tubulin, 1:1000), 133D3 (pS345-CHK1, 1:1000), rabbit polyclonal pT68-CHK2 (2661S, 1:1000) and pT183/Y185 p38MAPK mouse mAb 28B10 (1:2000) from Cell Signaling Technology (Danvers, MA); mouse mAbs 8E9 (total c-Abl, 1 μg/mL) and TU-01 (tubulin, 1:1000), and rabbit polyclonal Sdc2 (1 μg/mL) from Invitrogen/ThermoFisher Scientific (Rockford, IL); p53 mouse mAb DO-1 (1:750) from Santa Cruz Biotechnology (Dallas, TX) and mouse mAbs from Millipore-Sigma, JBW301 (pS139-γH2AX, 1:1000) and AC-74 (β-actin, 1:5000). Secondary antibodies for Western blotting include IRDye 680LT and 800CW goat anti-mouse IgG and goat anti-rabbit IgG (1:15,000, LI-COR Biosciences (Lincoln, NE), Alexa546 goat anti-rabbit and anti-mouse IgGs (1:5000, Invitrogen/ThermoFisher Scientific), alkaline phosphatase (AP)-conjugated donkey anti-goat, anti-rabbit and anti-mouse IgGs (1:5000, Jackson Immunoresearch, West Grove, PA). Antibodies for indirect immunofluorescence (IF) include : PCNA rabbit mAb D3H8P (0.2 μg/mL, Cell Signaling Technology), polyclonal MCM2 antibody (0.5 μg/mL, Bethyl Laboratories, Montgomery, TX) and whole IgG (Jackson Immunoresearch) followed by an Alexa546 goat anti-rabbit IgG secondary. Normal goat serum (VWR, Radnor, PA) diluted to 10% v/v in 1x CMF-PBS (pH 7.4) was used for IF blocking.

EGFR kinase inhibitors (erlotinib and gefitinib), MST1R/RON inhibitor BMS-0777607 and p38MAPK inhibitors (BIRB-796 and Losmapimod) were obtained from Selleck Chemicals (Houston, TX). EGFR-inhibitory antibody cetuximab was kindly provided by Dr. Paul Harari (University of Wisconsin-Madison). SFK inhibitor (PP2) and its paired inactive analogue (PP3), as well as the MST1R/RON inhibitor CAS 913376-84-8 were from Millipore-Sigma. Abl kinase inhibitors (GNF5 and PPY-A) were acquired from R&D Systems. EdU (5-Ethynyl-2′-deoxyuridine) and Click-IT EdU-labelling reagents (1.3 mM THPTA/CuSO_4_ mix, 20 µM AF488 picolyl azide, and 2.5 mM ascorbic acid) were from Click Chemistry Tools (Scottsdale, AZ) and Millipore-Sigma (CuSO_4_ and ascorbic acid). DDR stimulators (actinomycin D and hydroxyurea) and propidium iodide (PI; used to measure DNA content) were obtained from Millipore-Sigma. Human recombinant EGF was from Peprotech (Rocky Hill, NJ). CellTiter-GLO for quantifying cell numbers in proliferation assays was from Promega (Madison, WI).

### Cell culture

All cells were cultured at 37°C and 92.5% air/7.5% CO_2_. HaCaT normal human epidermal keratinocytes were kindly provided by Dr. Peter LaCelle (Roberts Wesleyan College, NY). Human normal oral keratinocytes (NOKs) and normal human tonsillary epithelial cells (HTEs) transduced with or without HPV oncogenes were kindly provided by Dr. Paul Lambert (UW-Madison). MCF10A immortalized human mammary epithelial cells, triple-negative human mammary carcinoma cells MDA-MB-231 and MDA-MB-468, SKBr3 (HER2+) human mammary carcinoma cells and SCC25 and UPCI-SCC90 human HNC cells were obtained from ATCC (Manassas, VA). UM-SCC47, UM-SCC1, and UM-SCC6 HNC cells were from Millipore-Sigma. TU-138 and UD-SCC2 HNC were kindly provided by Dr. Randy Kimple (UW-Madison). All HNC cells were authenticated by STR profile through Genetica LabCorp (Burlington, NC). NOKs and HTEs were cultured in complete Keratinocyte Serum-Free (KSFM) medium containing 100 units/mL penicillin and 100 μg/mL streptomycin (Life Technologies). All other cell lines were cultured as described previously^6–8, 112^. All cell lines were routinely screened for mycoplasma through the UW Small Molecule Screening Facility (SMSF) within the UW Carbone Cancer Center (UWCCC) Drug Development Core (DDC).

### Flow cytometry

To measure cell surface Sdc4, integrin and EGFR expression levels, suspended cells were incubated for 1 hr on ice with 1 µg of primary antibody per 3 × 10^5^ cells, washed, counterstained with Alexa-488-conjugated goat secondary antibodies and scanned on a ThermoFisher Attune NxT bench top cytometer.

Cell scatter and PI staining profiles were used to gate live, single-cell events for data analysis. For cell cycle analysis, asynchronous or synchronous (double thymidine block: 2 mM thymidine for 24 hr, 12 hr release in normal, complete culture medium followed by an additional 24 hr 2mM thymidine block) were cultured in the presence or absence of 30 µM SSTN_EGFR_ for 3, 8 (after release of the cells from the double thymidine block) or 16 hr. Cells were labelled with 100 μM EdU for the last 45 min of each SSTN incubation period prior to the cells being suspended, fixed in 4% EM-grade paraformaldehyde (PFA, ThermoFisher Scientific) and permeabilized in a buffer containing 1x CMF-PBS (pH 7.4), 0.5% saponin and 1% BSA (Millipore-Sigma). Cells were first stained using a Click-IT EdU-labelling reaction for 1 hr (1.3 mM THPTA/CuSO_4_ mix, 20 µM AF488 Picolyl Azide and 2.5 mM Ascorbic Acid in permeabilization buffer) followed by PI staining for 1.5 hr (20 μg/mL in the presence of 200 μg/mL DNAse-free RNase A (ThermoFisher Scientific) in permeabilization buffer). Cells were then analyzed by flow cytometry to assess levels of active DNA synthesis (AF488-EdU staining; Ex: 488 nm, Em: 530/30) and DNA content (PI staining; Ex: 561 nm, Em: 586/15) on a BD Biosciences LSR Fortessa bench top cytometer. Flow cytometry data acquisition and analysis were conducted at the UWCCC Flow Cytometry Laboratory Core.

### Immunoprecipitations and Western Blotting

Immunoprecipitation of the Sdc-RTK:ITG complex in the presence or absence of competing SSTN_EGFR_ peptide was carried out using 10 μg/mL Sdc4 mAb F94-8G3 (or Mouse IgG1as an isotype-matched, negative control), GammaBind PLUS Sepharose (GE Healthcare Life Sciences, Piscataway, NJ) and 1 mg of pre-cleared whole cell lysate (WCL)/sample in the presence of protease and phosphatase inhibitors as previously described^6, 8^. Immuno-isolated complexes were then enzymatically digested in 50 μL of heparitinase buffer (50 mM HEPES, 50 mM NaOAc, 150 mM NaCl, 5 mM CaCl2, pH 6.5) containing 4 × 10^-3^ IU/mL heparinase I and heparinase III (Syd Labs, Malden MA) and 0.1 conventional units/mL chondroitin ABC lysase (Millipore-Sigma) for 4 hr at 37°C (with fresh enzymes added after 2 hr) to remove the glycosaminoglycan (GAG) side chains, which is required to resolve the syndecan core proteins as a discrete bands by electrophoresis. To resolve Sdc4 from WCLs, 100 μg total protein was methanol-precipitated and then enzymatically digested with GAG lyases as described above. For p38MAPK and DDR effector blots, equal amounts of total protein (20-40 μg depending on the target protein and cell line being probed) - determined using the Pierce A660nm Protein Assay kit with Ionic Detergent Compatible Reagent (ThermoFisher Scientific) for SDS-PAGE sample buffer (lacking bromophenol blue) WCLs - were loaded per lane. All samples were resolved on 10% Laemmli gels prior to transfer to Immobilon-FL PVDF (Millipore-Sigma). Visualization of immunoreactive bands was performed using ECF reagent (for AP-conjugated secondary antibodies only) or direct excitation of fluorescent secondary antibodies on either a GE Healthcare Life Sciences Typhoon Trio (UWCCC Shared Equipment) or a LI-COR Odyssey Fc Imaging System (Department of Human Oncology, UW-Madison).

### siRNA design and transfection

*Silencer Select* control (AM4635) and target-specific siRNA oligos directed against human Sdc4 (siRNA ID# 12434; Target Sequence: ^197^(ca)GGAATCTGATGACTTTGAG^217^, GenBank Accession number NM_002999.2), ITGB4 (siRNA ID# s7584; Target Sequence: ^789^GCGACTACACTATTGGATT(tt)^807^, GenBank Accession Number NM_001005731.2), ITGA3 (siRNA ID# s7543; Target Sequence: ^1168^GGACTTATCTGAGTATAGT(tt)^1186^, GenBank Accession Number NM_002204.3) and EGFR (3’UTR Target Sequence: ^4899^GCTGCTCTGAAATCTCCTT^4917^, GenBank Accession Number NM_005228.4) were acquired from Life Technologies. UM-SCC47 cells (0.35 × 10^6^ per 35 mm well) were transfected with 100 nM siRNA using Lipofectamine RNAiMAX and Opti-MEM I transfection medium (from Life Technologies) at 1:1 ratio (μg siRNA:μL RNAiMAX). At 6 hr post-transfection, the wells were supplemented with 10% FBS and 3 mL of complete growth medium. At 24 hr post-transfection, the cells were suspended and plated on either acid-etched 18mm-#1 glass coverslips in 12-well plates or in 60mm tissue culture plates. Cells were then harvested at 72 h post-transfection and analyzed by Western blot (for receptor expression levels loading equal cell equivalents per sample) or AF488-EdU staining (for active DNA synthesis).

### Cell proliferation assays

Cells (1-2× 10^3^/well) were plated in 96-well plates in complete culture medium in the presence or absence of SSTN_EGFR_ peptide or EGFR kinase inhibitors for 72 hr. Cell numbers were measured using CellTiter-GLO against a standard curve in accordance with the manufacturer’s instructions.

### Transwell Filter Invasion Assays

The bottom polycarbonate filter surface of Transwell inserts (8 μm pores; Corning #3422) were coated with 10 μg/ml of LN332 (Kerafast, Boston, MA) diluted in 1x CMF-PBS (pH 7.4) for 3 hr at 37°C. NOKs or UM-SCC47 cells (5 × 10^4^) suspended in serum-free medium containing 1% heat-denatured BSA were plated in the upper insert chamber with or without 10 ng/ml EGFR and indicated inhibitors (Figs. 6B and C). Cells were allow to migrate/invade for 16 hr at 37°C in the tissue-culture incubator. Cells on the bottom of the filter were then fixed with 4% PFA and stained with 0.1% Crystal Violet (in ddH2O). Five random fields from duplicate inserts were imaged and quantified for each treatment condition.

### Immunofluorescence

Cells were plated on acid-etched 18mm-#1 glass coverslips in 12-well plates overnight and then treated with or without SSTN_EGFR_ peptide in the presence or absence of the indicated inhibitors (Figs. 6D, 7A and B, 8E and G) for 1, 3, or 6 hr. For EdU labelling, cells were incubated with 100 μM EdU in the last 45 mins of the treatment incubation period. The cells were fixed in 4% PFA, permeabilized in 0.5% Triton X-100 (Promega) in 1x CMF-PBS (pH 7.4) and blocked for 1 hr at room temperature in a 3% BSA/CMF-PBS solution. The cells were then stained using a Click-IT EdU-labelling reaction for 30 mins at room temperature (1.3 mM THPTA/CuSO_4_ mix, 20 µM AF488 Picolyl Azide and 2.5 mM Ascorbic Acid in a 1% BSA/CMF-PBS solution) prior to washing with PBT solution (CMF-PBS containing 1% BSA and 0.2% Tween-20) and mounting of the coverslips on glass slides using Prolong Diamond Mountant with DAPI (Life Technologies). For PCNA staining, cells were first hypotonically lysed in ice-cold ddH2O for 4 sec and then further permeabilized (to release non-DNA bound PCNA from the nucleus) in a lysis buffer containing 10 mM Tris (pH 7.4), 2.5 mM MgCl2, 0.5% NP-40, HALT protease and phosphatase inhibitor cocktail and 1 mM DTT for 10 min under constant agitation. The cells were then washed with 1x CMF-PBS (pH 7.4), blocked in PBT solution (also used for washes between staining steps) for 1 hr prior to Click-IT EdU AF488-labelling as described above. Cells were then stained with rabbit mAb D3H8P antibody (1:800) for 1.5 hr followed by Alexa546 goat anti-rabbit secondary antibody (1:500) for 45 mins prior to mounting. Fluorescent images (6 representative fields from duplicate wells/experiment) were acquired using either a Zeiss PlanAPOCHROMAT 10X (0.45 NA) or 20X objective (0.8 NA) and a Zeiss AxioCam Mrm CCD camera on a Zeiss Axio Imager.M2 microscopy system. EdU- and PCNA-positive stained cells were quantified using ImageJ (NIH, Bethesda, MD) and threshold limits set based on the relevant comparative control (i.e., untreated or solvent control-treated cells) for each experiment.

### SSTN Stability and PK/PD studies

Stability assays for SSTN_EGFR_ *in vitro* and *in vivo* (0.597 mg/kg) were conducted as previously described^8^.

### Tumor formation in animals

All animal studies were performed in accordance with UW and NIH guidelines after institutional review and approval. Under aseptic conditions, 2×10^6^ UM-SCC47 cells were subcutaneously injected into the right and left flanks of 6-8-wk-old female, athymic *Foxn1*^*nu*^ outbred nude mice (Envigo, East Millstone, NJ) in a 50:50 cell slurry in sterile PBS:Matrigel (BD Biosciences). Tumors were allowed to form for 1 week, then animals were randomized and surgically implanted with Alzet (Durect Corp., Cupertino, CA) osmotic pumps (Model 2004, 0.25 µl/h) containing either sterile saline or sterile saline plus 1.2 mM SSTN_EGFR_. After four weeks treatment, the mice were euthanized and tumors harvested and tumor volume (V = 0.524 × length × width^2^) determined^104^.

### Cell stress and apoptosis marker array

Screening for stress and/or apoptosis markers activated in cells treated with SSTN_EGF1R_ peptide for 16 hr (relative to vehicle-treated control cells) was conducted as previously described^8^ using the PathScan Stress and Apoptosis Signaling Antibody Array Kit from Cell Signaling Technology.

## ACKNOWLEDGMENTS

This work was supported by funds from the National Cancer Institute and National Institute of General Medical Sciences to A.C.R. (R01-CA163662,) P.F.L. (R35-CA 210807, P01-CA022443), and R.A.A. (R01-GM57549, R01-CA104708), funds to A.C.R. and P.F.L. from the UW Head and Neck Cancer SPORE (P50DE026787), funds in support of head and neck cancer research to A.C.R., P.F. L. and R.A.A. from the UW School of Medicine and Public Health and the UW Carbone Cancer Center, and the use of the UW Carbone Cancer Center shared services, supported by P30 CA014520.

## Authorship Contributions

D.M.B. participated in concept design, generation of data and writing the manuscript; K.S., N.S., S.E.N., D. L. and O.J. performed experiments and provided critique of the manuscript; R.A. A. and P.F.L. participated in concept design and critique of the manuscript, and A.C.R. oversaw concept design and participated in writing the manuscript.

